# Predicting Spatially Resolved Gene Expression via Tissue Morphology using Adaptive Spatial GNNs

**DOI:** 10.1101/2024.06.02.596505

**Authors:** Tianci Song, Eric Cosatto, Gaoyuan Wang, Rui Kuang, Mark Gerstein, Martin Renqiang Min, Jonathan Warrell

**Affiliations:** Department of Computer Science and Engineering, University of Minnesota, Minneapolis, 55455, MN, USA; Machine Learning Department, NEC Laboratories America, Princeton, 08540, NJ, USA; Program in Computational Biology and Bioinformatics, Yale University, New Haven, 06520, CT, USA; Department of Molecular Biophysics and Biochemistry, Yale University, New Haven, 06520, CT, USA; Department of Computer Science, Yale University, New Haven, 06520, CT, USA; Department of Statistics and Data Science, Yale University, New Haven, 06520, CT, USA; Department of Biomedical Informatics and Data Science, Yale University, New Haven, 06520, CT, USA

**Keywords:** Spatial Transcriptomics, Graph Neural Networks, Adaptive Graph, Spatial Gene Expression Prediction

## Abstract

**Motivation:** Spatial transcriptomics technologies, which generate a spatial map of gene activity, can deepen the understanding of tissue architecture and its molecular underpinnings in health and disease. However, the high cost makes these technologies difficult to use in practice. Histological images co-registered with targeted tissues are more affordable and routinely generated in many research and clinical studies. Hence, predicting spatial gene expression from the morphological clues embedded in tissue histological images, provides a scalable alternative approach to decoding tissue complexity.

**Results:** Here, we present a graph neural network based framework to predict the spatial expression of highly expressed genes from tissue histological images. Extensive experiments on two separate breast cancer data cohorts demonstrate that our method improves the prediction performance compared to the state-of-the-art, and that our model can be used to better delineate spatial domains of biological interest.

**Availability:** https://github.com/song0309/asGNN/

## Introduction

Dissecting the cellular and spatial heterogeneity of tissues is critical to characterizing cellular composition and organization, and ultimately their contribution to phenotype variation. Unlike traditional bulk and single-cell transcriptomics, spatial transcriptomics technologies enable spatially resolved gene expression profiling within intact tissues by using imaging or sequencing methods. Unlike imaging methods, which required targeted probes for a predetermined set of genes, sequencing-based methods, perform RNA sequencing of the whole transcriptome with a positionally barcoded array of spots aligned to the histological image of the tissue [1]. However, while sequencing-based spatial transcriptomics technologies have been widely used in biomedical research, their high cost still hinders their application in clinical studies. In contrast, histological images, such as hematoxylin and eosin (H&E) or immunofluorescence (IF) staining images, which are generated by most spatial transcriptomics technologies with ISC method, can be acquired cost-efficiently at high-quality. However, while these histological images are commonly used to compensate for spatial gene expression in downstream analyses [2], the dependencies between spatial gene expression profiles and histological images have only been explored to a limited extent; using such dependencies may alleviate the reliance on spatial transcriptomics by estimating spatial gene expression directly from tissue morphology.

As spatial transcriptomics data continues to accumulate, an increasing number of computational methods [4, 5, 6] aim to establish a connection between spatial gene expression profiles and histological images based on existing spatial transcriptomics datasets. These approaches predict the gene expression of each capturing spot with the corresponding image patch from H&E staining image. However, all of these methods fail to model spatial proximity in gene expression, which is one of the essential properties in real spatial transcriptomics data. A few attempts have been made to circumvent this issue, which either apply Transformer [7, 8] or GNN [9, 10] approaches to incorporate the relations among capturing spots when predicting spatial gene expression.

Despite the fact that transformer-based methods naturally model global relations among image patches and capturing spots by exploiting the self-attention mechanism, some methods attempt to further refine these relations by either incorporating positional information (e.g. HisToGene [7]) or imposing locality in the image embedding space (e.g. EGN [8]); they therefore lack the capability to distinguish compartments showing similar morphological features but distinct gene expressions in the tissues, such as tumors and their microenvironment. In addition, transformer-based methods may be prone to overfitting issue due to limited training data in the existing spatial transcriptomics datasets. Conversely, while GNN-based methods emphasize local relations among image patches and capturing spots in the graph, they may not retain global relations between spatially distant regions showing both similar morphological features and gene expression, particularly in the non-well-structured tissues, such as lymph nodes and tumors. Some methods allow both local and global relations to be encoded by considering both image and positional embeddings in the graph construction (e.g. SEPAL [9]), but these relations, which are hard-coded in the graph prior to model training, might not be present in gene expression.

To overcome the aforementioned issues, we propose an adaptive spatial Graph Neural Network (asGNN) for spatial gene expression prediction, which builds on the smoothing based GNN (SBGNN) framework of [3]. The SBGNN framework was developed to predict liquid-liquid phase separation from 3D molecular graphs, by using graph structure to adaptively refine molecular graphs to remove task irrelevant edges to help perform graph classification. Similarly, we adaptively remove edges in our spatial graph to help accurately predict gene expression. Following [3], during training we apply smoothing-based variational optimization [11] to search for a graph that captures both local and global relations important to a given task. In our case, these are relations among the capturing spots for a given image, and we use Graph Transformer Networks (GTN) [12] as the backbone to better align these relations with actual proximity in gene expression among capturing spots. Our experiments demonstrate that the spatial graphs learned from asGNN not only improve the spatial gene expression prediction compared to the state-of-the-art methods but also help to detect biologically interpretable spatial domains. Furthermore, the prototype clustering analysis on breast cancer tissues suggests that asGNN can be used to study the homogeneity and heterogeneity of spatial organization across tissue sections from patients in different conditions. The predictions from our model thus have the potential to be translated into clinical diagnostics tools to inform personalized treatment decisions.

## Methods

The overall architecture of our model is summarized in Figure 1. Below, we provide full details of the architecture and our end-to-end optimization algorithm, which build on the smoothing-based GNN framework of [3].

**Fig. 1.**
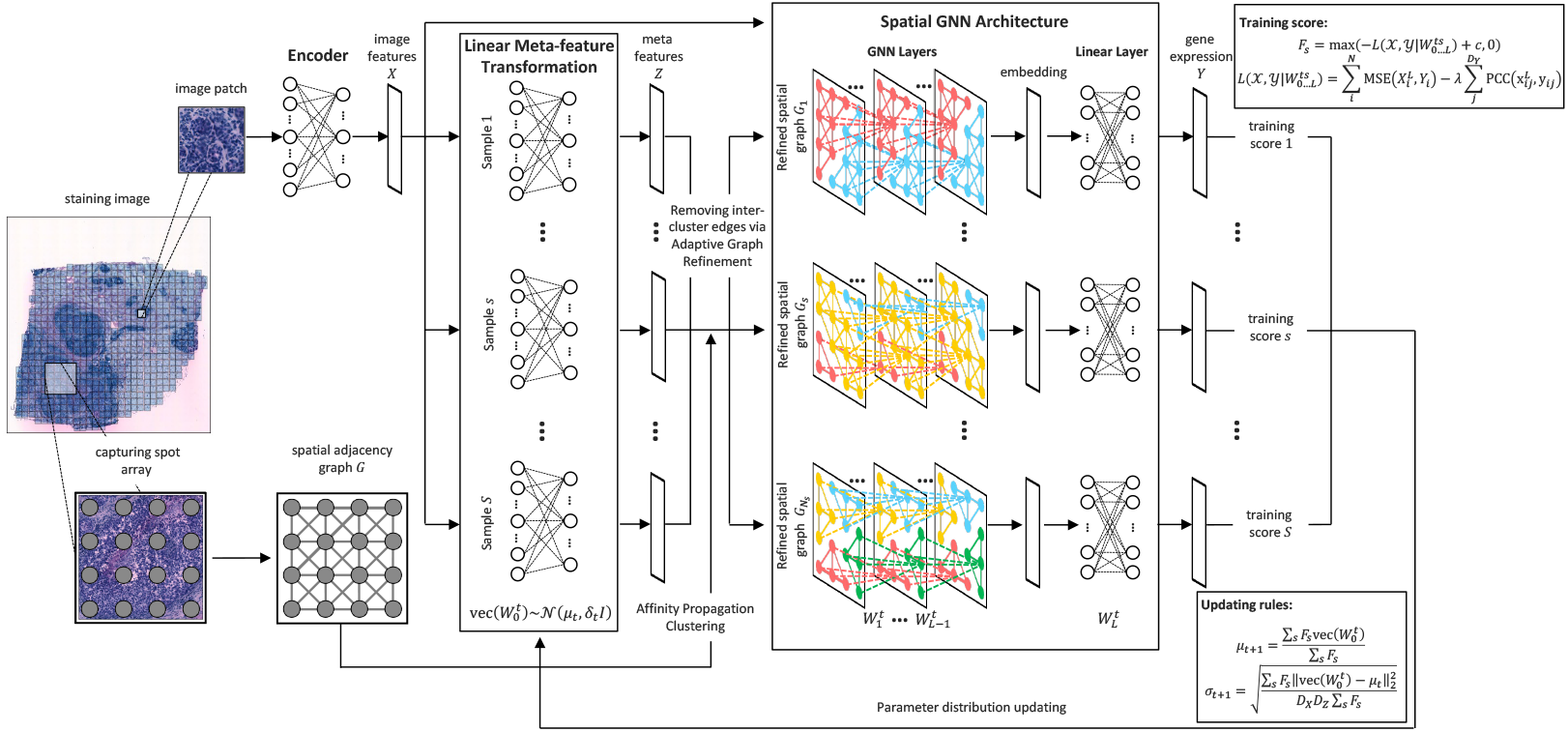
Adaptive Spatial Graph Neural Network (asGNN) Architecture. We summarize the key components of our asGNN architecture. asGNN begins by extracting image features from image patches over capturing spots arranged in an 8-connected spatial graph with an encoder. Adaptive graph refinement is then applied, as introduced in [3]; for each meta-epoch, asGNN samples *N*_*s*_ sets of parameters for a linear Meta-feature transformation, that projects the image features to a set of meta-features for each spot, and graph refinement is performed by applying Affinity Propagation (AP) clustering to the meta-features to sparsify the input graph. We use a GTN as the backbone model in asGNN, and train the ensemble of GTNs with image features on the sparsified spatial graphs to predict the gene expression separately. Lastly, we average the training scores over all the samples using the score function shown, which combines the mean squared error (MSE) and per-gene Pearson Correlation Coefficient (PCC) of the predicted gene-expression values, and apply variational updates to parameters of the linear Meta-feature transformation layer.

### Adaptive Spatial GNN Architecture

We assume we have input data of the form *χ* = {*G*_*i*_ = (*X*_*i*_, *E*_*i*_)|*i* = 1…*N* }, where *G*_*i*_ is the image graph for the *i*’th data point (a whole slide image), with *X*_*i*_ the matrix of node features for data point *i*, and *E*_*i*_ the edge set for the spatial connectivity of graph *G*_*i*_ (in our images, the spots are positioned in a regular grid, and we use an 8-connected neighborhood). *X*_*i*_ has dimensions *N*_*i*_ × *D*_*X*_, where *N*_*i*_ is the number of nodes (spots) in the image graph *G*_*i*_, and *D*_*X*_ is the dimensionality of the image features. Our task is then to predict the output spatial gene expression data, *Y* = {*Y*_*i*_|*i* = 1…*N* }, where *Y*_*i*_ is the expression matrix for image *i*, whose dimensionality is *N*_*i*_×*D*_*Y*_, where *D*_*Y*_ is the number of predicted genes.

#### Adaptive graph refinement

We adapt the graph refinement procedure introduced in [3]. During training, we learn to predict a matrix of latent meta-features, *Z*_*i*_ for each image, with dimensionality *N*_*i*_ × *D*_*Z*_, where *D*_*Z*_ is the meta-feature dimensionality. The latent meta-features in our model are learned as a linear transformation of the image features, i.e. *Z*_*i*_ = *X*_*i*_*W*_0_. We then define a distance function on the nodes of graph *i*:

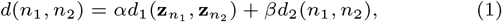

where **z**_*n*_ are the meta features associated with node *n* (i.e. the *n*’th row of *Z*_*i*_), *d*_1_ is the Euclidean distance, *d*_2_(*n*_1_, *n*_2_) is the shortest path distance between nodes *n*_1_ and *n*_2_ in *G*_*i*_, and {*α, β*} are hyperparameters. Our distance function here is adapted from [3], where we exclude their final ‘degree consistency’ term, due to the regular topology of our initial spatial graph. We use the distances in Eq. 1 to define a distance matrix *D*_*i*_ between each pair of nodes in graph *i*, and use an arbitrary clustering algorithm to map this to a vector *C*_*i*_ of cluster indices; here 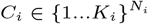, where *K*_*i*_ is the maximum cluster index for image *i*, and *C*_*i*_(*n*) = *k* indicates that node *n* belongs to cluster *k*. The clusters {1…*K*_*i*_} represent potentially meaningful spatial domains in image *i* (e.g. tumor micro-environment regions). We choose Affinity Propagation as our clustering algorithm [13], hence, *C*_*i*_ = AP(*D*_*i*_), where AP(.) denotes the application of the Affinity Propagation algorithm. Finally, using the learned cluster vectors, we form the refined spatial graphs for each training instance, 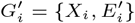, where 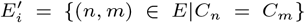, hence restricting the graph so that information is shared via message passing only within spatial domains (clusters).

#### Predicting spatial gene expression

We use the refined graphs to predict the spatial gene expression matrices as output; we thus adapt the framework of [3] (designed for graph classification tasks) to perform multivariate graph regression. Our network outputs the predicted matrix by performing message passing on the refined graphs, 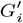. The network is parameterized by weight matrices *W*_1…*L*_, where *L* is the number of layers in the GNN, with *W*_*l*_ having dimensionality *D*_*l*−1_ × *D*_*l*_, such that *D*_*l*_ is the number of hidden units per node in layer *l*, and *D*_0_ = *D*_*X*_, *D*_*L*_ = *D*_*Y*_. We treat {*L, D*_1…*L*−1_} as additional hyperparameters. The message-passing updates can be written as:

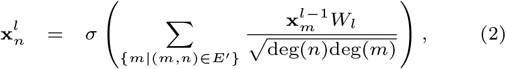

for levels *l < L*, where *σ*(*x*) = max(0, *x*) is the RELU function, and deg(*n*) is the degree of node *n*. For level *L*, a final linear is used, which is applied to each node independently; hence: 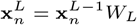

For our training loss, we use the function:

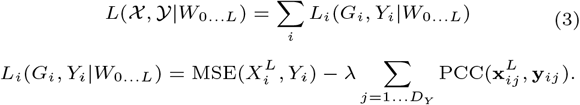

Here, MSE(*X, Y*) is the mean squared error between matrices *X* and *Y* (summed across all elements), PCC(**x, y**) is the Pearson Correlation Coefficient between vectors **x** and **y** (each being a vector of expression values across the nodes of the final layer), and *λ* is a trade-off parameter (which we set to 0 when considering the MSE loss only).

### End-to-end Training

Since, in our architecture, the projection *W*_0_ determines the meta-feature matrix *Z*, which in turn determines the graph structure of the adapted (refined) spatial graph *G*^*′*^ used for message passing, we have a complex interaction between a discrete optimization over the space of refined spatial graphs (implicitly parameterized by *W*_0_) and the continuous predictions of the network (determined by *W*_1…*L*_. The underlying objective (Eq. 3) is therefore discontinuous at points where changing *W*_0_ changes the refined graphs; however, if *W*_0_ is held constant, the objective is continuous over the remaining parameters, and can be handled by gradient descent.

Following [3], we thus use a modified form of Variational Optimization (VO) [11], which allows us to convert an objective with discontinuities into a continuous objective. This is done by introducing a variational distribution over the parameters *W*_0_, which we take to be a Gaussian with a symmetric covariance matrix. At a given meta-epoch *t*, this variational distribution has the form:

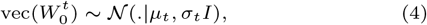

where vec(*W*_0_) is the vectorization of matrix *W*_0_, *μ*_*t*_ is a vector of mean values, *σ*_*t*_ is a scalar, and *I* is the identity matrix. At meta-epoch *t*, we draw *S* samples from Eq. 4, 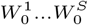, and for each we optimize the remaining parameters using gradient descent, to find 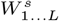 for *s* = 1…*S* (where we reserve a portion of the training data as a validation set to perform early stopping). Hence, we can calculate the sample training loss at meta-epoch *t*:

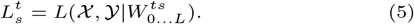

We then apply the smoothing-based optimization (SMO) updates from [11], to update *μ* and *σ*:

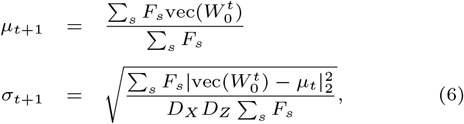

where *F*_*s*_ = max(−*L*_*s*_ + *c*, 0) is the score for sample *s*, with the offset *c >* 0 set to a positive constant (we treat *c* as an additional hyperparameter, which is set empirically to ensure that −*L*_*s*_ + *c >* 0 for observed values of *L*_*s*_). The updates in Eq. 6 can be shown to improve the value of *F* (i.e. the inverse loss) in expectation (as shown in [3, 11]); hence:

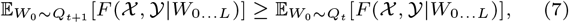

where *Q*_*t*_ = *𝒩*(.|*μ*_*t*_, *σ*_*t*_) is the variational distribution over *W*_0_ at meta-epoch *t*, and 𝔼[.] is the expectation operator.

## Experiments

### Data Preparation

In this work, we focus on spatial transcriptomics data generated by ST using the ST1K protocol [14], which measures the spatial expression of 26,949 genes by placing an array of 1,007 capturing spots arranged in a 33×35 grid onto the tissue section and provides a tissue image stained with hematoxylin and eosin (H&E). ST data were collected from two cohorts for breast cancer studies: One cohort [4] contains 68 tissue sections from 23 breast cancer patients with 5 different molecular subtypes, and the other cohort [15] consists of 36 tissue sections from 8 HER2-positive breast cancer patients. In the latter, each patient has one tissue section that was manually annotated with up to 5 tissue types based on the morphological features of the associated H&E staining image. We extracted image patches of 224 × 224 pixels centered on the corresponding capturing spots for each H&E staining image, where 224 × 224 pixels approximately cover each spot and is the standard input size for convolutional neural networks (CNNs) to derive a convenient feature set. For the spatial gene expression data, we followed the experimental setting in ST-Net [4], and first preprocessed the unique molecular identifier (UMI) counts in the raw data by normalizing them to sum to one after adding a pseudo count one for each capturing spot, and then transformed the normalized counts onto a log scale. Many of the genes are either lowly or sparsely expressed, and thus may not be essential for latent representation learning. Therefore, we followed the ST-Net setting [4] and filtered the top 250 genes with the highest gene expression across all tissue sections from two data cohorts for model training and prediction.

### Experimental Design

For spatial gene expression prediction, we benchmarked asGNN against 4 main baseline methods, including ST-Net [4], HisToGene [7], a basic (non-adaptive) GTN, and AP-Clustering+GTN (AP-GTN). The basic GTN was coupled with different spatial adjacency graphs by randomly dropping edges among capturing spots at ratios ranging from 0% (full) to 100% (empty), and the AP-GTN was combined with a spatial graph determined by AP clustering with similar distance function as Eq. (1) but defined on the untransformed image features. Note that other methods we mentioned in the introduction were excluded from the comparison since either their codes were not publicly available at the time of investigation or they showed unreasonably poor performance on the data in our experimental setting. We used two different image features, including morphological and convolutional features, with all the methods tested, except for ST-Net and HisToGene. The morphological features were calculated as a 142-dimensional vector concatenating morphological statistics and nuclei type proportion, derived from the nuclei segmentation produced by HoVerNet [16] on each image patch around a capturing spot. The convolutional features were calculated as a 1,024-dimensional vector extracted from pre-trained DenseNet-121 on ImageNet for each image patch around a capturing spot. Lastly, we assessed the performance of spatial gene expression prediction for all compared methods with both holdout and external validation sets. In the holdout validation, we stratified the 68 tissue sections from the first data cohort based on their molecular subtypes into training, validation, and test sets, consisting of 38, 15, and 15 sections, respectively, where the validation set was used to prevent overfitting in model training and select the best hyperparameters while the test set was used to evaluate the performance of the methods. In the external validation set, we employed 24 tissue sections from the second data cohort to evaluate the generalization of methods.

For spatial domain detection, we applied AP clustering to raw convolutional features and convolution-associated meta-features acquired from asGNN (*λ*=0) and asGNN for spot clustering separately. Without the need to specify the number of clusters, the performance of AP clustering may be degraded when an intact spatial domain is partitioned into separate regions. We thus first attempted to merge AP clusters by hierarchical clustering based on the averaged features per cluster, to find the best alignment of the detected spatial domains with the annotations. However, depending on the complexity of the tissue section, the clustering performance may still be suboptimal as delicate spatial domains can be merged with their surrounding domains. Therefore, we utilized the AP generated clusters to first sparsify the spatial adjacency graph, and then identified connected components (CC) in the sparsified graph as fine-grained clusters, before finally applying hierarchical clustering to merge these fine-grained clusters according to their averaged features to find the best alignment between the merged spatial domains and the annotations.

In the last experiment, we performed a prototype clustering analysis to investigate the spatial organization across tissue sections. We pooled spatial domains detected by the asGNN for all tissue sections and computed the average of the nuclei type proportions over the image patches within each spatial domain based on the nuclei segmentation results derived from HoVerNet. We then applied k-means to group spatial domains into either k=5 or k=10 prototype clusters according to their averaged nuclei type proportion. Furthermore, we used the Wilcoxon rank sum test to identify differentially expressed genes for each prototype cluster based on the corresponding spatial gene expression, and then performed Gene Ontology (GO) enrichment analysis on prototype clusters to explore their associated biological functions.

Finally, all the experiments were conducted on a cluster using 30 CPUs and 256GB RAM. In this environment, running the asGNN with *S* = 30 required roughly 1 hour of wall time on the training data for each meta-epoch. Despite the best-performing model typically being discovered in the earlier epochs, we ran 60 meta-epochs in total to ensure comprehensive exploration.

### Spatial Expression Prediction

To evaluate the spatial gene expression prediction performance, we initially applied asGNN, along with two state-of-the-art methods, ST-Net and HisToGene, for the holdout and external validations on the spatial transcriptomics data from 68 and 36 breast cancer tissue sections, respectively, where the validation performance was measured by two metrics, mean squared error (MSE) and Pearson correlation coefficient (PCC). It is evident that asGNN consistently achieved the best spatial gene expression prediction performance with the lowest MSE (0.696 and 0.932) and the highest PCC (0.113 and 0.312) across all tissue sections in both holdout and external validation sets, compared to the other baseline methods, as shown in Tables 1 and S1. ST-Net and HisToGene exhibit competitive prediction performance in terms of MSE (ST-Net: 0.712; HisToGene: 0.723) in the holdout validation, but demonstrate worse performance (ST-Net: 1.081; HisToGene: 1.297) in the external validation, which suggests these methods might be prone to overfitting, due to the limited training data in our experimental setting, and may be less generalizable to external data. The significance levels of improvement achieved by asGNN compared to baseline models are presented in Table S2; we note that a large majority of the comparisons (across diverse feature sets, test sets, performance metrics and training objectives), have high statistical significance, suggesting our approach provides a robust improvement over baseline methods.

**Table 1.** Spatial gene expression prediction performance comparison of state-of-the-art models ST-Net and HisToGene, basic GTN models with full (8-connected) spatial adjacency graphs, GTN model with sparsified spatial graph by AP clustering (AP-GTN), and the asGNN model, using both morphological and convolutional features from HoVerNet and DenseNet-121 as image features respectively, and performing both holdout and external validation on two data cohorts consisting of 68 and 32 breast cancer tissue sections separately. The basic GTN and AP-GTN models are as outlined in [3]. The best performance in terms of mean square error (MSE) and Pearson correlation coefficient (PCC) are underlined in each column of the table (See complete results in Table S1).

| Method | Spatial Graph | Morphological Features |  |  |  | Convolutional Features |  |  |  |
| --- | --- | --- | --- | --- | --- | --- | --- | --- | --- |
|  |  | Holdout |  | External |  | Holdout |  | External |  |
|  |  | *MSE* | *PCC* | *MSE* | *PCC* | *MSE* | *PCC* | *MSE* | *PCC* |
| ST-Net* | N/A | - | - | - | - | 0.712 | 0.065 | 1.081 | 0.292 |
| HisToGene* | N/A | - | - | - | - | 0.723 | 0.024 | 1.297 | 0.204 |
| GTN* | Full | 0.719 | 0.063 | 1.246 | 0.199 | 0.736 | 0.065 | 1.071 | 0.280 |
| AP-GTN* | Pre-clustered | 0.716 | 0.051 | 1.361 | 0.125 | 0.733 | 0.074 | 0.986 | 0.235 |
| asGNN* | Adaptive | **0.701** | 0.069 | 1.213 | 0.210 | 0.705 | 0.083 | 0.990 | 0.288 |
| GTN | Full | 0.710 | 0.090 | 1.240 | 0.204 | 0.711 | 0.101 | 0.961 | 0.297 |
| AP-GTN | Pre-clustered | 0.713 | 0.073 | 1.302 | 0.193 | 0.721 | 0.098 | 0.973 | 0.242 |
| asGNN | Adaptive | 0.703 | **0.103** | **1.208** | **0.212** | **0.696** | **0.113** | **0.932** | **0.312** |

\* denotes the models only optimize mean square error in their loss function.

To better understand how the adaptive spatial graph learned from asGNN informs spatial gene expression prediction, we introduced GTN models with varying levels of edge removal (dropout) applied to the full spatial graph as baseline methods; following the baseline comparisons used in [3]. The comparison between GTN models with different spatial adjacency graphs confirms that spatial gene expression prediction benefits from local relations among the capturing spots. The observation that asGNN outperforms all basic GTN models indicates that the local relations might be redundant and even misleading, and that pruning the spatial adjacency graph with guidance from the spatial gene expression prediction significantly improves the prediction performance. Further comparison between asGNN and AP-GTN suggests that hard-coded local relations are not always reflected in the actual spatial gene expression, resulting in overfitting to training data.

Furthermore, we assessed the spatial gene expression prediction performance of asGNN, AP-GTN, and GTN models with both morphological and convolutional features in both validation settings to explore the predictive power of different image features. Our results revealed that the performance improved significantly only when convolutional features were coupled with the asGNN, which indicates that while morphological features generally show high predictive performance, convolutional features have the potential to show higher concordance with spatial gene expression by establishing better local relations.

To investigate the importance of modeling gene correlation in spatial gene expression prediction, we conducted ablation studies by setting *λ* to 0 in the loss for asGNN and GTN models. The asGNN clearly shows better prediction performance compared to its variant with *λ*=0, which indicates that modeling gene correlation in GTNs improves the exploitation of local relations for spatial gene expression prediction.

Finally, we visualized the gene expression patterns generated by asGNN and the alternative methods (the baseline ST-Net and HisToGene) on breast tissue sections in the test set from the holdout validation, as shown in Figure 2. We selected the top predicted gene for each section from the asGNN, including COL1A2, MYL9, C4B, IGLL5, and GAS5, which are all spatially variable genes (assessed using SPARK [17], p-value*<* 0.05) and related to breast cancer. We then compared the spatial expression patterns of the selected genes with their ground-truth from associated spatial transcriptomics data and counterparts from ST-Net and HisToGene. It is clear that spatial expression patterns predicted by asGNN are highly correlated with their ground-truth patterns and exhibit appropriate continuity over neighboring spots, which demonstrates that asGNN is capable of capturing local relations in the spatial expression. Table S4 and Figures S7-9 further compare the top predicted genes by asGNN, ST-Net and HisToGene; as shown, there is only minimal overlap between the top genes predicted by each method dataset at tissue-specific levels, while out of the 15 selected tissue-specific genes (across Figures 2 and S8-9), asGNN performs best according to PCC on 9 of these (compared to 2 and 4 for ST-Net and HisToGene respectively).

**Fig. 2.**
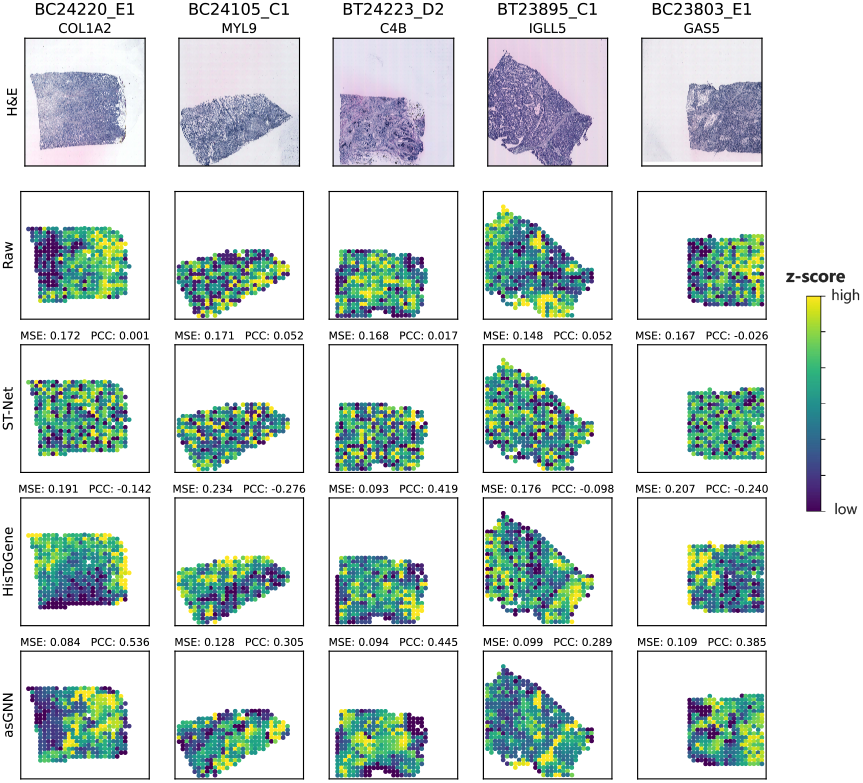
Expression pattern visualization of the top predicted genes by asGNN on 5 breast cancer tissue sections. Visualization of the raw and predicted expression patterns from ST-Net, HisToGene and asGNN for 5 spatially variable genes. Genes were selected by ranking prediction performance for each breast cancer tissue sections in the holdout validation set. Both mean squared error (MSE) and Pearson correlation coefficient (PCC) between raw and predicted expression are reported for each gene. Further visualizations of the top predicted genes by ST-Net and HisToGenes are shown in Figures S8 and S9.

### Spatial Domain Detection

One of the key advantages of asGNN is that it performs spot clustering implicitly in refining the spatial adjacency graph, where clusters obtained from either AP clustering on the latent meta-features or the sparsified spatial graph can be interpreted as spatial domains within the tissue section. To provide a quantitative measure of the spatial domains detected by asGNN, we evaluated spot clustering performance by computing the adjusted rand index (ARI) between spot clusters and annotations on 8 breast cancer tissue sections from the external validation set. ARIs were further optimized by merging either AP or CC clusters based on latent meta-features with hierarchical clustering, as described above in the Experimental Design. To better understand the importance of latent meta-features in spatial domain detection, we introduced a naive method in the comparison, which directly applied AP clustering to raw image features to generate AP and CC clusters. Note that we only focus on the convolutional features and their latent meta-features in this experiment, due to their superior performance in spatial gene expression prediction.

As shown in Figure S1, asGNN outperforms other baseline methods regardless of merging strategies and achieves the overall best ARI (median ARI = 0.423) by merging CC clusters for spatial domain detection, which indicates fine-grained CC clusters with convolution-associated meta-features learned from asGNN have the potential to accurately delineate spatial domains. We observed that convolution-associated meta-features generally improve the spot clustering performance compared to raw convolutional features in both merging strategies, and noticed a substantial improvement by merging CC clusters (p-value*<*0.05), which implies that convolution-associated meta-features are more informative in capturing location relations. Interestingly, asGNN shows slightly worse ARI (median ARI = 0.322) than those from asGNN (*λ*=0) (median ARI = 0.363) when merging AP clusters, which might be attributed to delicate spatial domains being masked by the coarse-grained clusters obtained from AP clustering, possibly as a result of using non-optimal clustering hyperparameters.

To illustrate how the CC merging strategy and convolution-associated meta-features contribute to spatial domain detection intuitively, we further visualized the spot clustering results on annotated breast cancer tissue sections, as depicted in Figure 3. We found that the spatial domains from asGNN matched well with the annotated tissue regions and even exhibited high agreement with fine-grained structures (e.g. breast gland and immune infiltrate), whereas the spatial domains from naive methods appeared over-smoothed and failed to distinguish spatial domains correctly. asGNN and asGNN (*λ*=0) demonstrated superior performance in spatial domain detection by merging AP clusters (as shown in Figure S2); asGNN could identify continuous spatial domains with clear boundaries and asGNN (*λ*=0) could even recognize narrow structures consisting of a few spots (e.g. immune infiltrate). We note, however, that asGNN tends to produce fewer AP clusters than the actual number of annotated regions for most tissues, which might obfuscate some fine-grained structures.

**Fig. 3.**
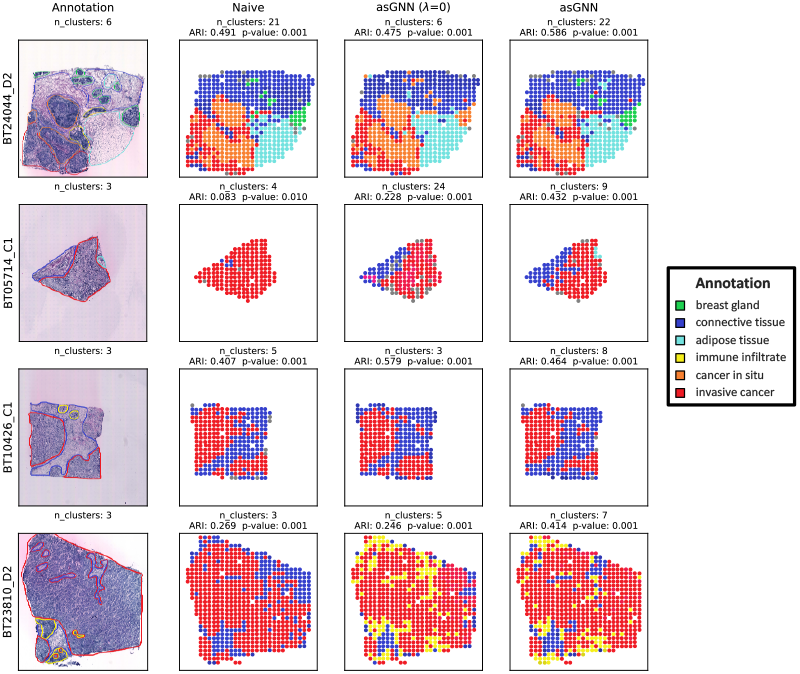
Spatial domain visualization on 4 breast cancer tissues. Visualization of spatial regions annotated by pathologists and spatial domains optimized by merging connected-component (CC) clusters from naive, asGNN (*λ*=0) and asGNN on 4 breast cancer tissue sections. Adjusted Rand Index (ARI) between detected spatial domains and annotated spatial regions, along with the corresponding p-values from permutation test, were reported for each method across different tissues. Note that the spatial domains are colored with different hues if they were recognized as the same region in the annotation. Singleton clusters were excluded for better visualization.

### Prototype Clustering and Enrichment Analysis

To further chart the spatial organization under various tissue contexts, we employed k-means clustering to discover prototypes for the spatial domains across all tissue sections from breast cancer patients. Instead of relying on latent meta-features, which might be biased towards spatial expression of particular genes, we used nuclei type composition derived from nuclei segmentation in the prototype clustering. As depicted in Figure 4, the spatial organization shows visual consistency across replicated tissue sections, implying the robustness of asGNN in spatial domain detection. While prototype clusters from different granularities of clustering exhibit high correspondence, the prototype clustering with k=10 allows finer-grained characterization spatial organization in breast cancer, such as tumor regions with different subtypes (clusters 0, 1, and 6).

**Fig. 4.**
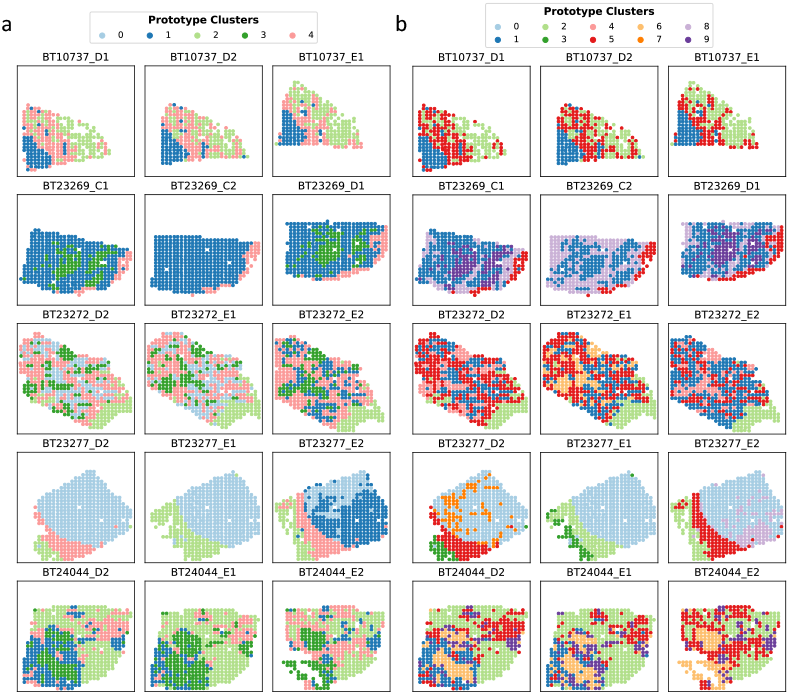
Prototype cluster visualization on 15 breast cancer tissue sections. Discovering prototype clusters across different breast cancer tissue sections by applying k-means to nuclei type composition of their AP clusters from the asGNN method (nuclei type composition determined using the nuclei segmentation results from HoVerNet). **a** Visualization of prototype clusters (*k* = 5). **b** Visualization of prototype clusters (*k* = 10). Further examples are shown in Figures S5 and S6.

To evaluate the stability of the cluster prototypes, we measured the Stability Score (SS) for each prototype cluster by calculating the bootstrapped ARIs of the clustering results after masking out each prototype cluster in turn. Subsequently, to further investigate the biological interpretation of prototype clusters, we conducted GO enrichment analysis on the differentially expressed genes from all the genes in the original spatial gene expression data (not only the predicted genes) for each prototype cluster for both k=5 and k=10 clustering settings, and the top enriched GO terms along with the stability scores for each prototype cluster are present in Table S3. We identified prototype clusters with SS*>*0.7 and found their enriched biological processes to be highly relevant to the corresponding regions in the annotated tissue sections; for example, clusters 1 and 6 in the k=10 setting are enriched for tumor development and tumor-associated immune responses [18], respectively, closely resembling the tumor core and surrounding regions in a few tissue sections. Interestingly, we also discovered that several prototype clusters, which are enriched for biological processes related to the microenvironment, are spatially adjacent to each other in some breast cancer tissue sections, which aligns well with the concept of a multilayered microenvironment [19].

## Discussion

We have introduced an adaptive spatial Graph Neural Network architecture (asGNN) for spatial gene expression prediction, which builds on the adaptive graph refinement framework of [3]. We have shown that our model generates state-of-the-art performance for predicting spatial gene expression from histological image data. Our method learns to adapt the spatial graph structure on an image-by-image basis, so that information is only shared between spots in coherent spatial domains, defined by the learned meta-features. Our model can be trained in an end-to-end fashion, using a smoothing-based variational optimization approach [3, 11]. Further, we have shown that the spatial domains identified by our method achieve a high degree of alignment with pathologist annotations, and can be readily interpreted biologically through our prototype analysis as corresponding to layered tumor and tumor microenviroment regions.

As future work, we intend to investigate both the biological and clinical potential of our method. The ready availability of large quantities of histology images (for instance, in TCGA [20]) suggests that we may be able to improve the predictive performance or fine-tune our model to different tumors using a pseudo-labeling (semi-supervised) approach, by augmenting our training set with instances where only image or image and bulk expression data are available. Further, the ability of our approach to generate putative spatial domains, suggests that we may be able to identify *de novo* tumor-type specific tumor or microenvironment domains through such analysis. We further plan to test the potential of our approach to handle spatial expression data with higher spatial resolution (for instance, subcellular), and explore the potential for using cluster identity to influence graph refinement (to model inter-cluster dependencies). Finally, we intend to investigate the potential of our method to identify spatial biomarkers for patient stratification for clinical diagnostics and personalized treatment, where the spatial expression patterns predicted by our model can be used both as biomarkers themselves, and, in a semi-supervised setting, to help learn novel biomarkers. Code available at: https://github.com/song0309/asGNN/

## Acknowledgements

This work was supported by NEC Laboratories America.

## References

1. Michaela Asp, Joseph Bergenstråhle, and Joakim Lundeberg. Spatially resolved transcriptomes—next generation tools for tissue exploration. BioEssays, 42(10):1900221, 2020.

2. Ruben Dries, Jiaji Chen, Natalie Del Rossi, et al. Advances in spatial transcriptomic data analysis. Genome research, 31(10):1706–1718, 2021.

3. Gaoyuan Wang, Jonathan Warrell, Suchen Zheng, et al. A variational graph partitioning approach to modeling protein liquid-liquid phase separation. bioRxiv preprint doi: 10.1101/2024.01.20.576375, 2024.

4. Bryan He, Ludvig Bergenstråhle, Linnea Stenbeck, et al. Integrating spatial gene expression and breast tumour morphology via deep learning. engineering, 4(8):827–834, 2020. Nature biomedical

5. Taku Monjo, Masaru Koido, Satoi Nagasawa, et al. Efficient prediction of a spatial transcriptomics profile better characterizes breast cancer tissue sections without costly experimentation. Scientific Reports, 12(1):4133, 2022.

6. Muhammad Dawood, Kim Branson, Nasir M Rajpoot, et al. All you need is color: image based spatial gene expression prediction using neural stain learning. In Joint European Conference on Machine Learning and Knowledge Discovery in Databases, pages 437–450. Springer, 2021.

7. Minxing Pang, Kenong Su, and Mingyao Li. Leveraging information in spatial transcriptomics to predict super-resolution gene expression from histology images in tumors. bioRxiv, pages 2021–11, 2021.

8. Yan Yang, Md Zakir Hossain, Eric A Stone, et al. Exemplar guided deep neural network for spatial transcriptomics analysis of gene expression prediction. In Proceedings of the IEEE/CVF Winter Conference on Applications of Computer Vision, pages 5039–5048, 2023.

9. Gabriel Mejia, Paula Cárdenas, Daniela Ruiz, et al. Sepal: Spatial gene expression prediction from local graphs. In Proceedings of the IEEE/CVF International Conference on Computer Vision, pages 2294–2303, 2023.

10. Yuansong Zeng, Zhuoyi Wei, Weijiang Yu, et al. Spatial transcriptomics prediction from histology jointly through transformer and graph neural networks. Briefings in Bioinformatics, 23(5):bbac297, 2022.

11. Marius Leordeanu and Martial Hebert. Smoothing-based optimization. In 2008 IEEE Conference on Computer Vision and Pattern Recognition, pages 1–8. IEEE, 2008.

12. Yunsheng Shi, Zhengjie Huang, Shikun Feng, et al. Masked label prediction: Unified message passing model for semi-supervised classification. arXiv preprint 2009.03509, 2020.

13. Brendan Frey and Delbert Dueck. Clustering by passing messages between data points. Science, 315(5814), 2007.

14. Sheel Shah, Eric Lubeck, Wen Zhou, et al. In situ transcription profiling of single cells reveals spatial organization of cells in the mouse hippocampus. Neuron, 92(2):342–357, 2016.

15. Alma Andersson, Ludvig Larsson, Linnea Stenbeck, et al. Spatial deconvolution of her2-positive breast cancer delineates tumor-associated cell type interactions. Nature communications, 12(1):6012, 2021.

16. Eric Cosatto, Pierre-Francois Laquerre, Christopher Malon, et al. Automated gastric cancer diagnosis on h&e-stained sections; ltraining a classifier on a large scale with multiple instance machine learning. In Medical Imaging 2013: Digital Pathology, volume 8676, pages 51–59. SPIE, 2013.

17. Shiquan Sun, Jiaqiang Zhu, and Xiang Zhou. Statistical analysis of spatial expression patterns for spatially resolved transcriptomic studies. Nat. methods, 17(2):193–200, 2020.

18. Karin E de Visser and Johanna A Joyce. The evolving tumor microenvironment: From cancer initiation to metastatic outgrowth. Cancer Cell, 41(3):374–403, 2023.

19. Lucie Laplane, Dorothée Duluc, Nicolas Larmonier, et al. The multiple layers of the tumor environment. Trends in cancer, 4(12):802–809, 2018.

20. John N Weinstein, Eric A Collisson, Gordon B Mills, et al. The cancer genome atlas pan-cancer analysis project. Nature genetics, 45(10):1113–1120, 2013.

