## Supplementary figures and tables for "Predicting Spatially Resolved Gene Expression via Tissue Morphology using Adaptive Spatial GNNs"

### Abstract

**Motivation:** Spatial transcriptomics technologies, which generate a spatial map of gene activity, can deepen the understanding of tissue architecture and its molecular underpinnings in health and disease. However, the high cost makes these technologies difficult to use in practice. Histological images co-registered with targeted tissues are more affordable and routinely generated in many research and clinical studies. Hence, predicting spatial gene expression from the morphological clues embedded in tissue histological images, provides a scalable alternative approach to decoding tissue complexity.
**Results:** Here, we present a graph neural network based framework to predict the spatial expression of highly expressed genes from tissue histological images. Extensive experiments on two separate breast cancer data cohorts demonstrate that our method improves the prediction performance compared to the state-of-the-art, and that our model can be used to better delineate spatial domains of biological interest.
**Availability:** https://github.com/song0309/asGNN/

**Key words:** Spatial Transcriptomics, Graph Neural Networks, Adaptive Graph, Spatial Gene Expression Prediction

### Related Works

**ST-Net** [1] fine-tunes a DenseNet-121 encoder followed by a fully connected layer to predict the gene expression of each capturing spot with the corresponding image patch from an H&E staining image.
**NSL** [2] learns a deconvolutional layer to estimate stain intensities for the image patch of each capturing spot to predict the corresponding gene expression.
**HisToGene** [3] projects image patches and spot coordinates into image and position embeddings with a learnable linear layer, and adopts a Vision Transformer (ViT) encoder on the aggregated embeddings of both image and positions to predict spatial gene expression.
**EGN** [4] trains a Generative Adversarial Network (GAN) model StyleGAN on image patches centered on capturing spots for reconstruction, where the encoder is then used to extract the embeddings for those image patches, and identifies the neighborhoods of the image patches in the embedding space to further guide a ViT to predict spatial gene expression.
**SEPAL** [5] first fine-tunes a ViT encoder followed by a linear layer on image patches to predict the gene expression of corresponding capturing spots, and constructs a similarity graph among spots by their image embeddings from the ViT encoder and spatial coordinates, and then applies a Graph Convolutional Network (GCN) to predict spatial gene expression.
**Hist2ST** [6] first utilizes a ConvMixer encoder to extract embeddings from image patches centered on capturing spots and further fuses spatial coordinates into the image embeddings with a Transformer encoder, and defines the spatial graph among spots based on their spatial arrangement. The method then applies a GCN model GraphSAGE to fit spatial gene expression with zero-inflated negative binomial (ZINB) distribution.

Note that we were only able to compare asGNN with ST-Net and HisToGene, since the code for the other methods above either were not publicly available at the time of investigation or they showed unreasonably poor performance on the data in our experimental setting (the latter was the case for: NSL, EGN, and SEPAL).

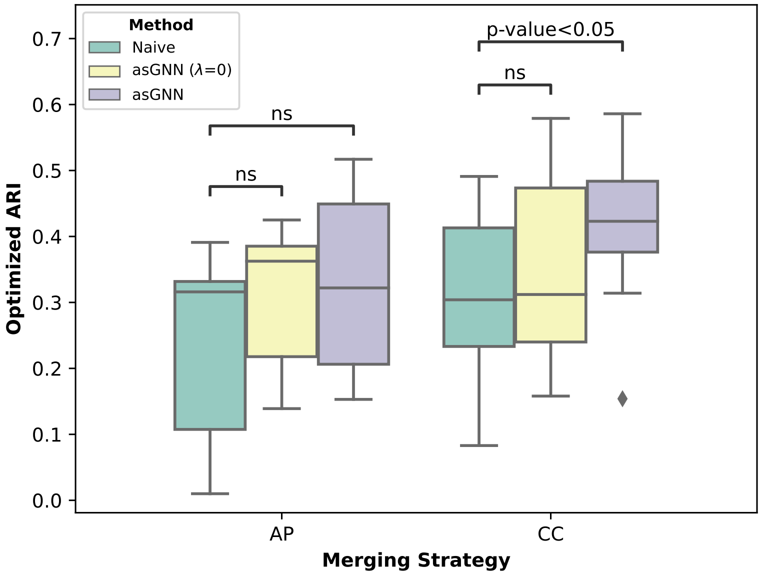


**Fig. S1. Spatial domain detection performance comparison between two merging strategies on clusters from different methods** The Adjusted Rand Index (ARI) on all 8 annotated breast cancer tissue sections was optimized by either merging AP clusters obtained from AP clustering or connected-component (CC) clusters derived from the sparsified spatial graph for each section, and both merging strategies were tested on clusters generated by naive, asGNN and asGNN ($\lambda = 0$) models. Paired Student t-tests were performed between the naive and asGNN methods for each merging strategy, and ns represents not significant. ARI optimization curves using AP and CC merging strategies are shown as Figures S3 and S4, respectively.

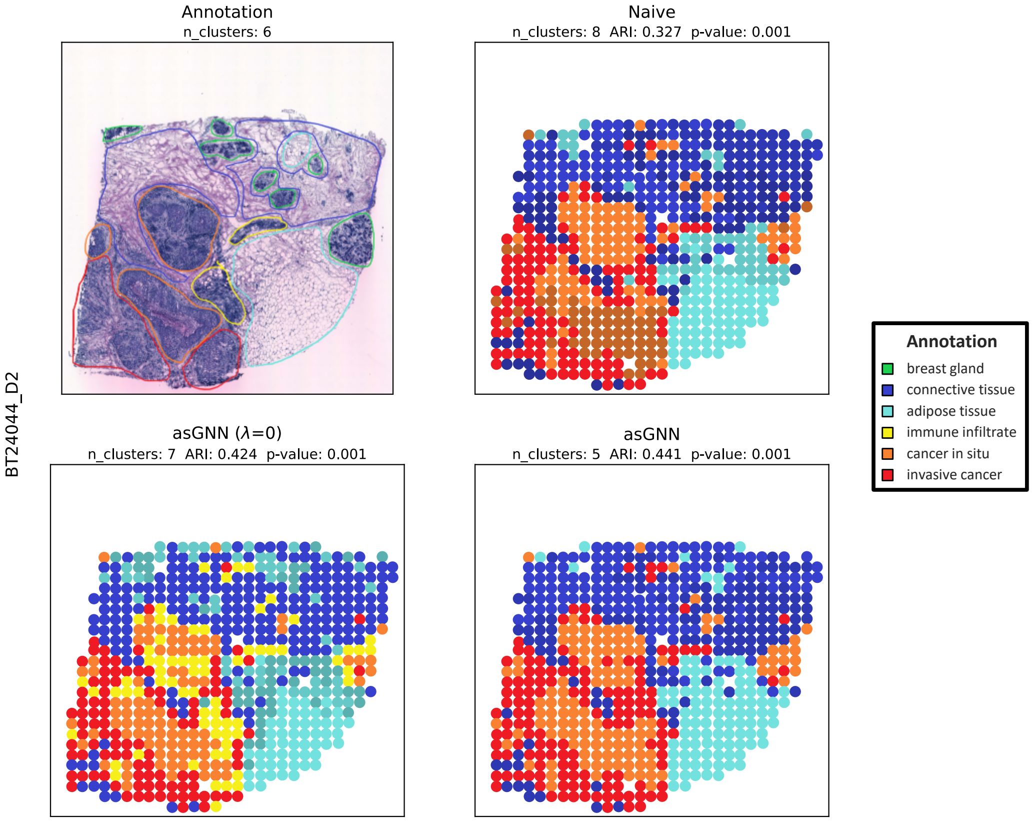


**Fig. S2. Spatial domain visualization by merging AP clusters**. Visualization of spatial regions annotated by pathologists and spatial domains detected by merging AP clusters from naive, asGNN ($\lambda$=0) and asGNN models on the tissue from the patient BT24044. Adjusted Rand Index (ARI) between detected spatial domains and annotated spatial regions, along with the corresponding p-values from the permutation test, were reported for each method. Similar to Figure 3, spatial domains are colored with different hues if they were recognized as the same region in the annotation, and singleton clusters were excluded as well.

**Table S1.** Spatial gene expression prediction performance comparison of state-of-the-art models ST-Net and HisToGene, basic GTN models with different spatial adjacency graphs, GTN model with sparsified spatial graph by AP clustering (AP-GTN), and the asGNN model, using both morphological and convolutional features from HoVerNet and DenseNet-121 as image features respectively, and performing both holdout and external validation on two data cohorts consisting of 68 and 32 breast cancer tissue sections separately. The best performance in terms of mean square error (MSE) and Pearson correlation coefficient (PCC) are underlined in each column of the table. Note that the performance of all methods below exhibited relatively large variances (calculated across the images in the test set) due to the highly diverse molecular subtypes present in both the holdout and external datasets. However, significant improvements are observed compared to all baseline methods for many conditions when the performance is compared across matching images via paired Student t-test comparisons; see Table S2 for a full summary of the statistical test results.

| Method | Spatial Graph | Morphological Features | | | | Convolutional Features | | | |
|---|---|---|---|---|---|---|---|---|---|
| | | Holdout | | External | | Holdout | | External | |
| | | *MSE* | *PCC* | *MSE* | *PCC* | *MSE* | *PCC* | *MSE* | *PCC* |
| **ST-Net*** | N/A | - | - | - | - | 0.712 ± 0.133 | 0.065 ± 0.143 | 1.081 ± 0.277 | 0.292 ± 0.145 |
| **HisToGene*** | N/A | - | - | - | - | 0.723 ± 0.172 | 0.024 ± 0.125 | 1.297 ± 0.359 | 0.204 ± 0.112 |
| **GTN*** | Empty | 0.718 ± 0.127 | 0.053 ± 0.073 | 1.285 ± 0.335 | 0.186 ± 0.110 | 0.739 ± 0.160 | 0.036 ± 0.102 | 1.010 ± 0.216 | 0.262 ± 0.143 |
| | Full | 0.719 ± 0.123 | 0.063 ± 0.083 | 1.246 ± 0.284 | 0.199 ± 0.128 | 0.736 ± 0.156 | 0.065 ± 0.083 | 1.071 ± 0.252 | 0.280 ± 0.175 |
| | Random (drop 25%) | 0.714 ± 0.168 | 0.053 ± 0.073 | 1.474 ± 0.290 | 0.097 ± 0.054 | 0.722 ± 0.166 | 0.078 ± 0.077 | 1.041 ± 0.290 | 0.226 ± 0.165 |
| | Random (drop 50%) | 0.713 ± 0.164 | 0.048 ± 0.068 | 1.475 ± 0.379 | 0.107 ± 0.063 | 0.727 ± 0.155 | 0.072 ± 0.086 | 1.065 ± 0.208 | 0.259 ± 0.190 |
| | Random (drop 75%) | 0.716 ± 0.150 | 0.064 ± 0.066 | 2.158 ± 0.212 | 0.048 ± 0.043 | 0.725 ± 0.170 | 0.079 ± 0.074 | 1.577 ± 0.308 | 0.137 ± 0.100 |
| **AP-GTN*** | Pre-clustered | 0.716 ± 0.160 | 0.051 ± 0.051 | 1.361 ± 0.370 | 0.125 ± 0.098 | 0.733 ± 0.139 | 0.074 ± 0.071 | 0.986 ± 0.202 | 0.235 ± 0.127 |
| **asGNN*** | Adaptive | 0.701 ± 0.130 | 0.069 ± 0.070 | 1.213 ± 0.327 | 0.210 ± 0.120 | 0.705 ± 0.111 | 0.083 ± 0.086 | 0.990 ± 0.226 | 0.288 ± 0.162 |
| **GTN** | Empty | 0.712 ± 0.130 | 0.067 ± 0.080 | 1.263 ± 0.332 | 0.195 ± 0.110 | 0.713 ± 0.144 | 0.094 ± 0.098 | 0.999 ± 0.239 | 0.295 ± 0.138 |
| | Full | 0.710 ± 0.128 | 0.090 ± 0.090 | 1.240 ± 0.311 | 0.204 ± 0.129 | 0.711 ± 0.133 | 0.101 ± 0.093 | 0.961 ± 0.202 | 0.297 ± 0.177 |
| | Random (drop 25%) | 0.710 ± 0.142 | 0.077 ± 0.083 | 1.250 ± 0.261 | 0.149 ± 0.084 | 0.714 ± 0.152 | 0.091 ± 0.105 | 0.968 ± 0.241 | 0.284 ± 0.167 |
| | Random (drop 50%) | 0.716 ± 0.146 | 0.076 ± 0.072 | 1.373 ± 0.411 | 0.111 ± 0.063 | 0.715 ± 0.134 | 0.089 ± 0.081 | 1.065 ± 0.254 | 0.266 ± 0.159 |
| | Random (drop 75%) | 0.704 ± 0.141 | 0.097 ± 0.085 | 1.940 ± 0.237 | 0.057 ± 0.045 | 0.716 ± 0.159 | 0.087 ± 0.092 | 1.313 ± 0.288 | 0.146 ± 0.091 |
| **AP-GTN** | Pre-clustered | 0.713 ± 0.133 | 0.073 ± 0.081 | 1.302 ± 0.347 | 0.193 ± 0.125 | 0.721 ± 0.137 | 0.098 ± 0.070 | 0.973 ± 0.200 | 0.242 ± 0.123 |
| **asGNN** | Adaptive | 0.703 ± 0.133 | 0.103 ± 0.096 | 1.208 ± 0.335 | 0.212 ± 0.126 | 0.696 ± 0.126 | 0.113 ± 0.101 | 0.932 ± 0.187 | 0.312 ± 0.163 |

∗ denotes the models only optimize mean square error in their loss function.

**Table S2.** Significance of prediction performance improvement for asGNN models compared with the state-of-the-art models ST-Net and HisToGene, basic GTN models with full spatial adjacency graphs, as well as GTN model with sparsified spatial graph by AP clustering (AP-GTN). Paired Student t-tests were performed on the mean square error (MSE) and Pearson correlation coefficient (PCC) between asGNN and baseline models. The significant improvements with p-values $< 0.05$ are underlined in each column of the table.

| Method | Spatial Graph | Morphological Features | | | | Convolutional Features | | | |
|---|---|---|---|---|---|---|---|---|---|
| | | Holdout | | External | | Holdout | | External | |
| | | *MSE* | *PCC* | *MSE* | *PCC* | *MSE* | *PCC* | *MSE* | *PCC* |
| **ST-Net*** | N/A | - | - | - | - | 8.86e-2[1]<br>4.57e-2[2] | 9.27e-2[1]<br>5.00e-2[2] | 6.24e-5[1]<br>1.07e-6[2] | 7.34e-1[1]<br>4.51e-2[2] |
| **HisToGene*** | N/A | - | - | - | - | 7.48e-2[1]<br>2.72e-2[2] | 1.31e-2[1]<br>5.95e-4[2] | 2.71e-9[1]<br>2.85e-9[2] | 2.66e-5[1]<br>4.78e-7[2] |
| **GTN*** | Full | 5.03e-3[1] | 3.94e-1[1] | 1.28e-2[1] | 3.45e-6[1] | 4.65e-2[1] | 1.34e-1[1] | 1.01e-4[1] | 5.51e-1[1] |
| **AP-GTN*** | Pre-clustered | 2.50e-2[1] | 2.66e-1[1] | 9.39e-12[1] | 3.45e-6[1] | 1.07e-4[1] | 5.05e-1[1] | 7.83e-1[1] | 3.96e-4[1] |
| **asGNN*** | Adaptive | 3.46e-1[2] | 2.04e-2[2] | 5.63e-1[2] | 8.64e-1[2] | 8.86e-2[2] | 1.29e-2[2] | 1.62e-3[2] | 3.96e-2[2] |
| **GTN** | Full | 4.51e-2[2] | 5.60e-1[2] | 1.07e-3[2] | 5.47e-1[2] | 1.08e-3[2] | 9.68e-2[2] | 4.80e-2[2] | 4.75e-3[2] |
| **AP-GTN** | Pre-clustered | 4.24e-2[2] | 2.76e-2[2] | 2.07e-8[2] | 1.08e-1[2] | 2.66e-3[2] | 2.62e-3[2] | 3.72e-2[2] | 4.53e-5[2] |
| **asGNN** | Adaptive | - | - | - | - | - | - | - | - |

* denotes the models only optimize mean square error in their loss function.

[1] indicates the results are compared with asGNN*.

[2] indicates the results are compared with asGNN.

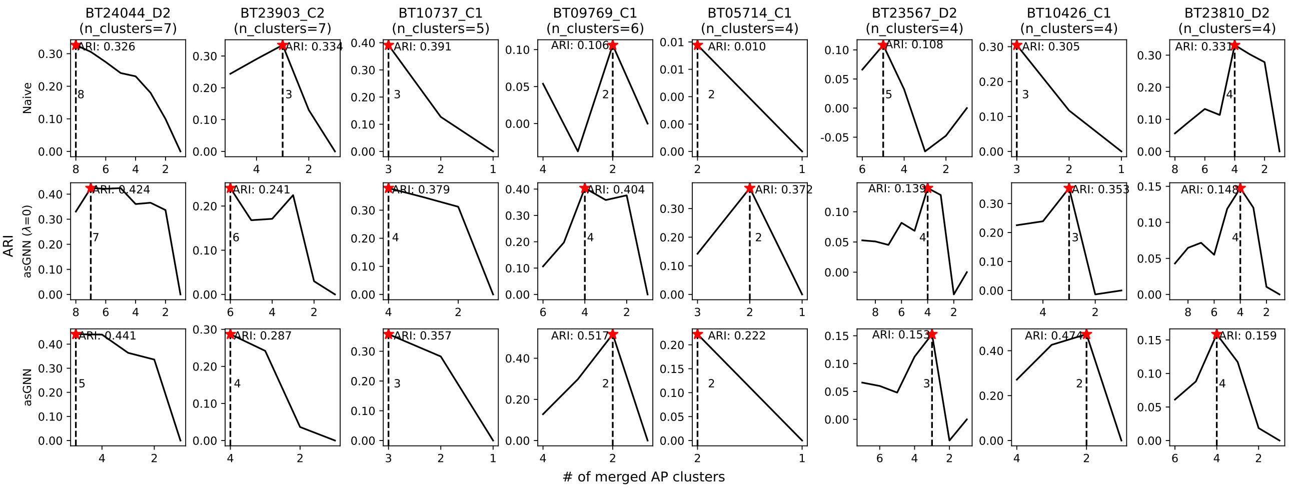


**Fig. S3. Spatial domain optimization using AP-cluster merging strategies**. ARI curves between spatial regions and AP clusters were calculated for naive, asGNN ($\lambda$=0), and asGNN after each merging, using hierarchical clustering on 8 annotated breast cancer tissue sections. The best ARI and the corresponding number of merged AP clusters are reported for each method across different tissues.

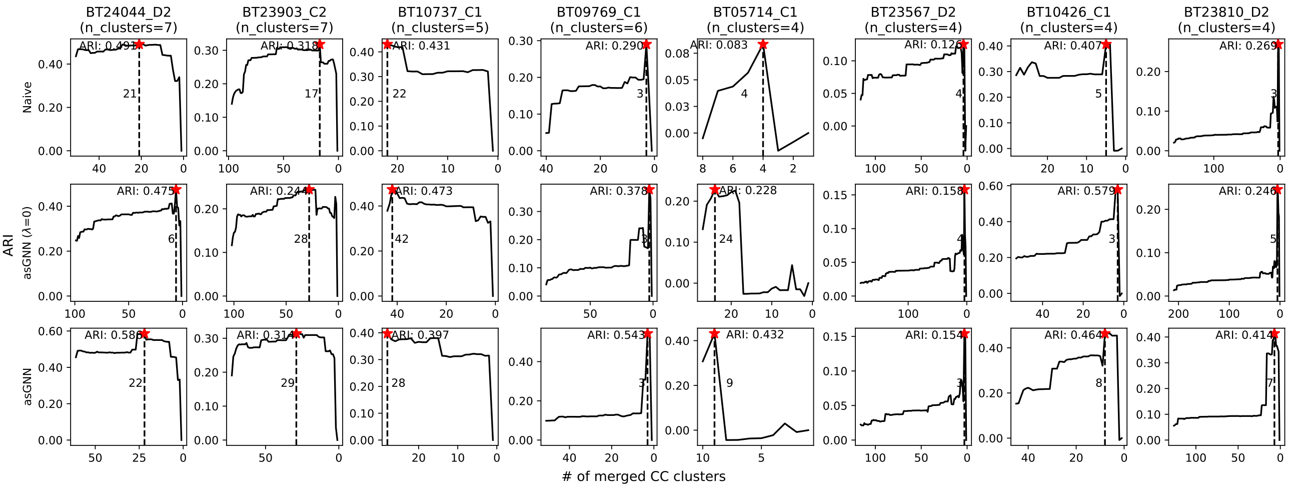


**Fig. S4. Spatial domain optimization using CC-cluster merging strategies**. ARI curves between spatial regions and connected-component (CC) clusters (i.e. splitting each AP cluster into connected components on the spatial graph) were calculated for naive, asGNN ($\lambda$=0), and asGNN after each merging, using hierarchical clustering on 8 annotated breast cancer tissue sections. The best ARI and the corresponding number of merged CC clusters are reported for each method across different tissues.

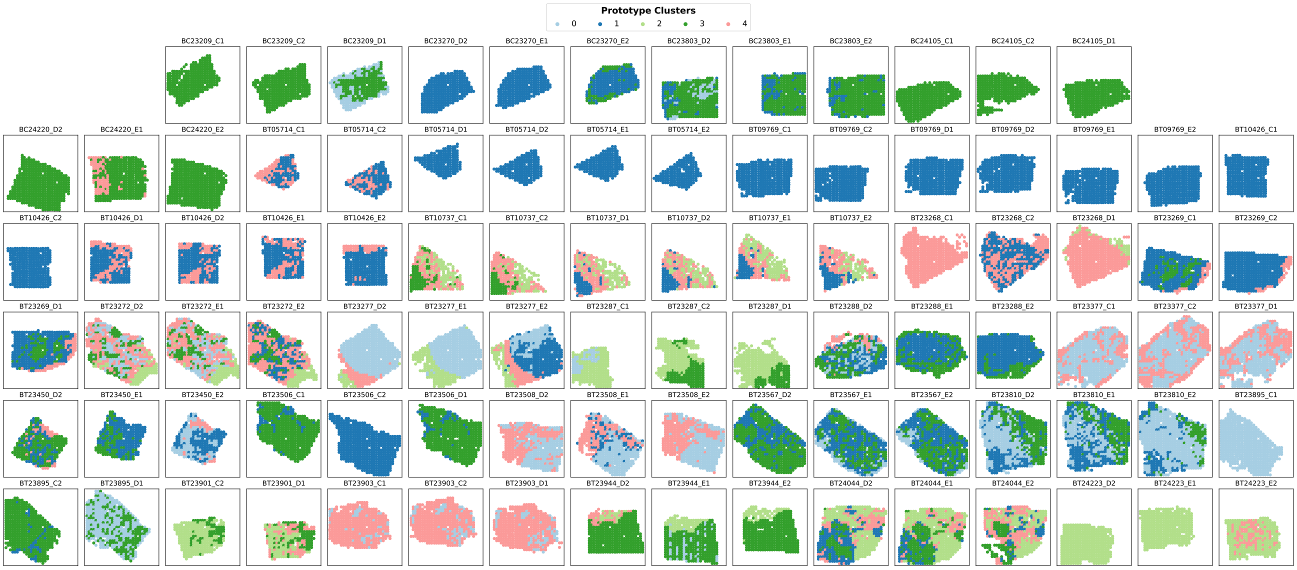


**Fig. S5. Prototype cluster visualization on 92 breast cancer tissue sections** Discovering prototype clusters across different breast cancer tissue sections by applying k-means (k=5) to nuclei type composition of the AP clusters associated with the asGNN method (nuclei type composition determined using the nuclei segmentation results from HoVerNet).

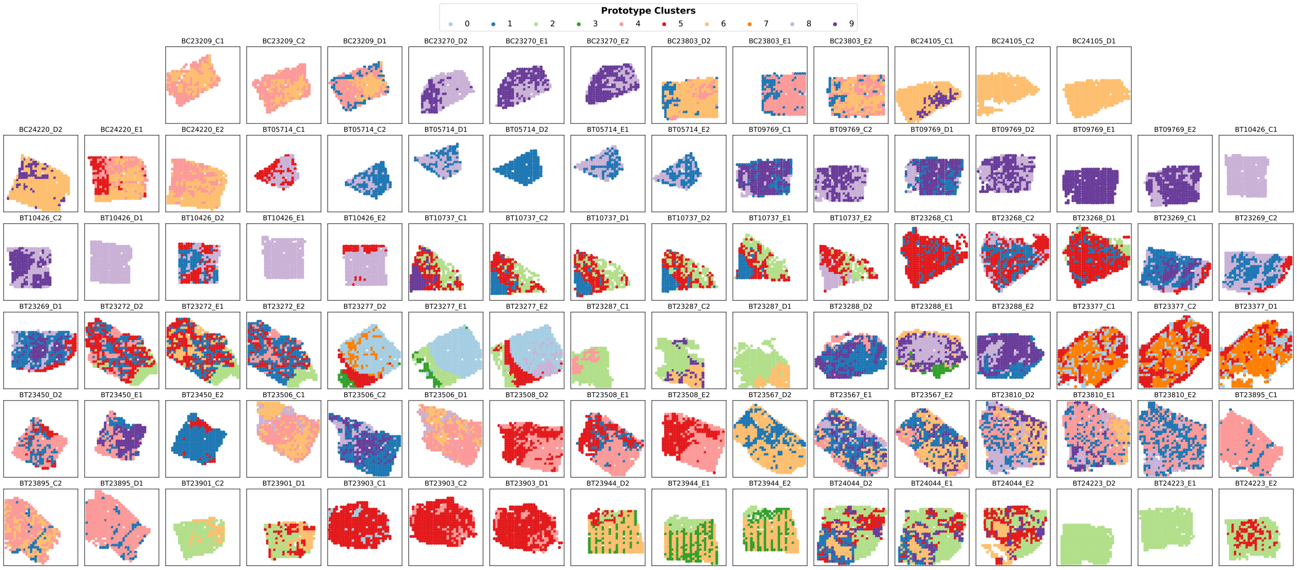


**Fig. S6. Prototype cluster visualization on 92 breast cancer tissue sections** Discovering prototype clusters across different breast cancer tissue sections by applying k-means (k=10) to nuclei type composition of the AP clusters associated with the asGNN method (nuclei type composition determined using the nuclei segmentation results from HoVerNet).

**Table S3.** Enrichment analysis on prototype clusters across 92 breast cancer tissue sections. Top 4 enriched GO terms, along with their corresponding p-values are reported for each prototype clusters. In addition, the stability score was calculated for each cluster (see text) in both prototype clustering settings (k=5 and k=10). The clusters with stability scores exceeding 0.7 are underlined in the table.

| # of Prototype Clusters | Cluster ID | Stability Score | Top Enriched Gene Ontology (GO) Terms (p-values) | Summary |
|---|---|---|---|---|
| **k = 5** | 0 | 0.444 | Phosphorylation (GO:0016310) (9.03e-3)<br>Cytosolic Transport (GO:0016482) (3.15e-2)<br>Regulation Of Gluconeogenesis (GO:0006111) (3.26e-2)<br>Mitochondrial Electron Transport, Ubiquinol To Cytochrome C (GO:0006122) (3.70e-2) | Tumor* |
| | 1 | 0.553 | Intracellular Protein Transmembrane Transport (GO:0065002) (1.90e-2)<br>Protein N-linked Glycosylation Via Asparagine (GO:0018279) (1.90e-2)<br>Mitochondrial Translation (GO:0032543) (2.56e-2)<br>Endoplasmic Reticulum To Golgi Vesicle-Mediated Transport (GO:0006888) (4.82e-2) | Tumor Metastasis* |
| | 2 | 0.555 | Aerobic Electron Transport Chain (GO:0019646) (1.27e-4)<br>Mitochondrial ATP Synthesis Coupled Electron Transport (GO:0042775) (1.27e-4)<br>NADH Dehydrogenase Complex Assembly (GO:0010257) (1.90e-4)<br>Oxidative Phosphorylation (GO:0006119) (3.94e-3) | TME |
| | 3 | <u>0.727</u> | Interleukin-27-Mediated Signaling Pathway (GO:0070106) (4.35e-3)<br>Cholesterol Biosynthetic Process (GO:0006695) (7.23e-3)<br>Type I Interferon-Mediated Signaling Pathway (GO:0060337) (7.23e-3)<br>Sterol Biosynthetic Process (GO:0016126) (1.07e-2) | Tumor Inflammation |
| | 4 | <u>0.716</u> | Regulation Of Angiogenesis (GO:0045765) (2.61e-13)<br>Regulation Of Cell Migration (GO:0030334) (3.64e-10)<br>Extracellular Matrix Assembly (GO:0085029) (1.21e-9)<br>Elastic Fiber Assembly (GO:0048251) (7.51e-9) | TME |
| **k = 10** | 0 | 0.407 | Mitochondrial Electron Transport, Ubiquinol To Cytochrome C (GO:0006122) (1.51e-2)<br>Cellular Response To Decreased Oxygen Levels (GO:0036294) (1.51e-2)<br>Cellular Response To Hypoxia (GO:0071456) (2.34e-2)<br>Positive Regulation Of Mitochondrion Organization (GO:0010822) (2.34e-2) | Tumor |
| | 1 | 0.600 | Positive Regulation Of Phagocytosis (GO:0050766) (1.89e-3)<br>Immunoglobulin Mediated Immune Response (GO:0016064) (4.78e-3)<br>Positive Regulation Of Leukocyte Cell-Cell Adhesion (GO:1903039) (5.61e-3)<br>Positive Regulation Of Endocytosis (8.86e-3) | Tumor Immune Infiltration |
| | 2 | <u>0.793</u> | Aerobic Electron Transport Chain (GO:0019646) (9.08e-4)<br>Mitochondrial ATP Synthesis Coupled Electron Transport (GO:0042775) (1.89e-3)<br>Regulation Of Protein Kinase B Signaling (GO:0051896) (5.61e-3)<br>Positive Regulation Of Cell Motility (GO:2000147) (5.61e-3) | TME |
| | 3 | 0.493 | Cytokine-Mediated Signaling Pathway (GO:0019221) (7.07e-3)<br>Regulation Of I-kappaB kinase/NF-kappaB Signaling (GO:0043122) (1.68e-2)<br>Response To Type II Interferon (GO:0034341) (2.11e-2)<br>Interleukin-27-Mediated Signaling Pathway (GO:0070106) (2.12e-2) | TME |
| | 4 | 0.598 | Ubiquitin-Dependent Protein Catabolic Process (8.52e-5)<br>Positive Regulation Of Proteasomal Protein Catabolic Process (GO:1901800) (3.03e-3)<br>Regulation Of mRNA Stability (GO:0043488) (5.59e-3)<br>Protein Phosphorylation (GO:0006468) (7.59e-3) | Tumor* |
| | 5 | <u>0.785</u> | Regulation Of Angiogenesis (GO:0045765) (7.02e-10)<br>Regulation Of Cell Migration (GO:0030334) (1.22e-9)<br>Extracellular Matrix Assembly (GO:0085029) (1.22e-9)<br>Elastic Fiber Assembly (GO:0048251) (1.50e-7) | TME |
| | 6 | <u>0.716</u> | Cholesterol Biosynthetic Process (GO:0006695) (1.26e-4)<br>Sterol Biosynthetic Process (GO:0016126) (2.25e-4)<br>Positive Regulation Of Inflammatory Response (GO:0050729) (3.19e-2)<br>Interleukin-27-Mediated Signaling Pathway (GO:0070106) (3.19e-2) | Tumor Inflammation |
| | 7 | 0.631 | Regulation Of DNA-templated Transcription (GO:0006355) (5.07e-9)<br>Regulation Of Transcription By RNA Polymerase II (GO:0006357) (6.32e-6)<br>Regulation Of mRNA Splicing, Via Spliceosome (GO:0048024) (5.19e-3)<br>Transmembrane Receptor Protein Tyrosine Kinase Signaling Pathway (GO:0007169) (1.70e-2) | Tumor* |
| | 8 | <u>0.776</u> | Protein N-linked Glycosylation Via Asparagine (GO:0018279) (2.30e-3)<br>Regulation Of Cell Communication By Electrical Coupling (GO:0010649) (4.52e-2)<br>Positive Regulation Of Apoptotic Process (GO:0043065) (4.52e-2)<br>Cellular Response To Interferon-Beta (GO:0035458) (4.52e-2) | Tumor Metastasis* |
| | 9 | 0.636 | DNA Repair (GO:0006281) (3.34e-9)<br>DNA-templated DNA Replication (GO:0006261) (6.91e-7)<br>Intrinsic Apoptotic Signaling Pathway (GO:0097193) (1.25e-3)<br>Positive Regulation Of Cell Cycle Process (GO:0090068) (2.32e-3) | Tumor* |

∗ denotes the summary of the prototype cluster is less confident based on its enrichment results. TME denotes tumor microenvironment

**Table S4.** Top 10 genes predicted by ST-Net, HisToGene, and asGNN. The genes were ranked by their average Pearson Correlation Coefficient (PCC) against the ground-truth expression over 15 breast cancer tissue sections in the holdout validation set. The intersection of the top-performing genes across all three methods is highlighted in the table.

| Method | Performance Ranking | | | | | | | | | |
|---|---|---|---|---|---|---|---|---|---|---|
| **ST-Net*** | PSMD3 | RPL13 | EVL | RPL29 | H2AJ | CCND1 | CD81 | KRT18 | **ACTG1** | H1-10 |
| **HisToGene*** | **ACTG1** | ATP1A1 | DDX5 | EVL | FASN | GNAS | H3-3B | HSP90AB1 | PTMA | XBP1 |
| **asGNN** | COL1A1 | **ACTG1** | AEBP1 | FN1 | COL1A2 | GNAS | IGLL5 | FASN | SPARC | TMSB10 |

* denotes the models only optimize mean square error in their loss function.

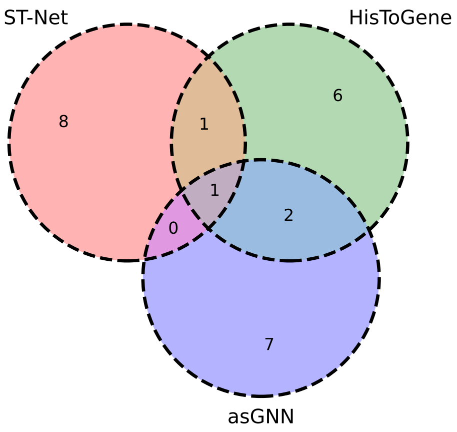


**Fig. S7. Overlap of top 10 predicted genes by different methods.** Venn diagram of overlaps between the top 10 predicted genes from ST-Net, HisToGene and asGNN. Genes were selected by ranking average prediction performance for each method over 15 breast cancer tissue sections in the holdout validation set.

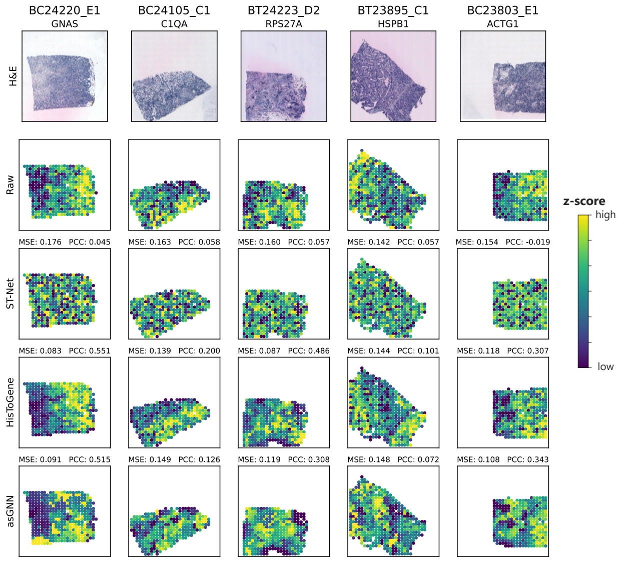


**Fig. S9. Expression pattern visualization of the top predicted genes by HisToGene on 5 breast cancer tissue sections.** Visualization of the raw and predicted expression patterns from ST-Net, HisToGene and asGNN for 5 spatially variable genes. Genes were selected by ranking prediction performance of HisToGene for each breast cancer tissue section in the holdout validation set. Both mean squared error (MSE) and Pearson correlation coefficient (PCC) between raw and predicted expression are reported for each gene.

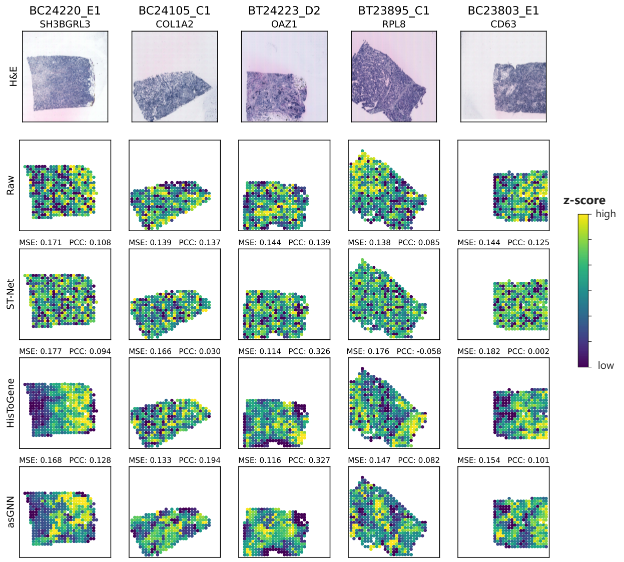


**Fig. S8. Expression pattern visualization of the top predicted genes by ST-Net on 5 breast cancer tissue sections.** Visualization of the raw and predicted expression patterns from ST-Net, HisToGene and asGNN for 5 spatially variable genes. Genes were selected by ranking prediction performance of ST-Net for each breast cancer tissue section in the holdout validation set. Both mean squared error (MSE) and Pearson correlation coefficient (PCC) between raw and predicted expression are reported for each gene.

### References

1. Bryan He, Ludvig Bergenstråhle, Linnea Stenbeck, et al. Integrating spatial gene expression and breast tumour morphology via deep learning. *Nature biomedical engineering*, 4(8):827–834, 2020.
2. Muhammad Dawood, Kim Branson, Nasir M Rajpoot, et al. All you need is color: image based spatial gene expression prediction using neural stain learning. In *Joint European Conference on Machine Learning and Knowledge Discovery in Databases*, pages 437–450. Springer, 2021.
3. Minxing Pang, Kenong Su, and Mingyao Li. Leveraging information in spatial transcriptomics to predict super-resolution gene expression from histology images in tumors. *bioRxiv*, pages 2021–11, 2021.
4. Yan Yang, Md Zakir Hossain, Eric A Stone, et al. Exemplar guided deep neural network for spatial transcriptomics analysis of gene expression prediction. In *Proceedings of the IEEE/CVF Winter Conference on Applications of Computer Vision*, pages 5039–5048, 2023.
5. Gabriel Mejia, Paula Cárdenas, Daniela Ruiz, et al. Sepal: Spatial gene expression prediction from local graphs. In *Proceedings of the IEEE/CVF International Conference on Computer Vision*, pages 2294–2303, 2023.
6. Yuansong Zeng, Zhuoyi Wei, Weijiang Yu, et al. Spatial transcriptomics prediction from histology jointly through transformer and graph neural networks. *Briefings in Bioinformatics*, 23(5):bbac297, 2022.
